# *Cis*-regulatory changes associated with a recent mating system shift and floral adaptation in *Capsella*

**DOI:** 10.1101/017749

**Authors:** Kim A. Steige, Johan Reimegård, Daniel Koenig, Douglas G. Scofield, Tanja Slotte

## Abstract

The selfing syndrome constitutes a suite of floral and reproductive trait changes that have evolved repeatedly across many evolutionary lineages in response to the shift to selfing. Convergent evolution of the selfing syndrome suggests that these changes are adaptive, yet our understanding of the detailed molecular genetic basis of the selfing syndrome remains limited. Here, we investigate the role of *cis-*regulatory changes during the recent evolution of the selfing syndrome in *Capsella rubella*, which split from the outcrosser *Capsella grandiflora* less than 200 kya. We assess allele-specific expression (ASE) in leaves and flower buds at a total of 18,452 genes in three interspecific F1 *C. grandiflora* x *C. rubella* hybrids. Using a hierarchical Bayesian approach that accounts for technical variation using genomic reads, we find evidence for extensive *cis-*regulatory changes. On average, 44% of the assayed genes show evidence of ASE, however only 6% show strong allelic expression biases. Flower buds, but not leaves, show an enrichment of *cis-*regulatory changes in genomic regions responsible for floral and reproductive trait divergence between *C. rubella* and *C. grandiflora*. We further detected an excess of heterozygous transposable element (TE) insertions near genes with ASE, and TE insertions targeted by uniquely mapping 24-nt small RNAs were associated with reduced expression of nearby genes. Our results suggest that *cis*-regulatory changes have been important during the recent adaptive floral evolution in *Capsella* and that differences in TE dynamics between selfing and outcrossing species could be important for rapid regulatory divergence in association with mating system shifts.

## Introduction

The transition from outcrossing to predominant self-fertilization has occurred repeatedly in flowering plants (Stebbins 1950). In association with this shift, marked changes in floral and reproductive traits have occurred independently in many different lineages (Barrett 2002). In general, selfers tend to show reduced allocation of resources to traits involved in pollinator attraction and reward (e.g. smaller petals, less nectar per flower, less scent), exhibit changes in floral morphology that may improve the efficacy of autonomous self-pollination (e.g. reduced separation between stigma and anthers), and show reduced allocation of resources to male function (reduced ratio of pollen to ovules) (reviewed in Sicard and Lenhard 2011). Together, this combination of floral and reproductive traits is termed “the selfing syndrome” (Ornduff 1969).

Despite the striking pattern of convergent floral evolution in association with the shift to selfing, we currently have a limited understanding of the molecular genetic basis of the selfing syndrome. Quantitative trait loci (QTL) for the selfing syndrome have been identified in a handful of systems (e.g. *Capsella*; Sicard et al 2011; Slotte et al 2012; *Leptosiphon*; Goodwillie et al 2006; *Mimulus*; Fishman et al 2002; Fishman et al 2015; *Oryza*; Grillo et al 2009; *Solanum*; Bernacchi and Tanksley 1997). In domesticated tomatoes, *cis-*regulatory changes at the *Style2.1* gene have been implicated in reduced stigma exsertion (Chen et al 2007), but in most other systems, the molecular basis of the selfing syndrome is not known. A major unresolved question thus concerns the general importance of *cis-*regulatory changes vs. other types of molecular changes for the evolution of the selfing syndrome.

*Cis-*regulatory changes have long been hypothesized to be important for organismal adaptation (Doebley and Lukens 1998; Carroll 2000; Wray 2007; Carroll 2008; Stern and Orgogozo 2008; but see Hoekstra and Coyne 2007), due to their potentially limited negative pleiotropic effects (Wray 2007). The prospects for identifying *cis*-regulatory changes on a transcriptome-wide scale have greatly improved due to the advent of massively parallel sequencing (Fraser 2011). In particular, methods for assessing allele-specific expression (ASE) that contrast the relative levels of expression of two alleles in an individual allow for transcriptome-scale assessment of *cis*-regulatory changes. ASE studies require the presence of transcribed polymorphisms as well as rigorous bioinformatic approaches, but have benefits over mapping approaches (e.g. eQTL mapping) in terms of cost and resolution, and can identify individual genes with *cis*-regulatory changes (Pastinen 2010).

As part of our broad goal to examine molecular genetic changes associated with the selfing syndrome, we examine the influence of *cis*-regulatory changes on the evolution of the selfing syndrome in *Capsella rubella*. We further test whether silencing of TEs through the RNA-directed methylation pathway is important for global *cis-*regulatory divergence in association with the shift to selfing. The crucifer genus *Capsella* is a promising system for assessing the role of *cis-*regulatory changes in association with plant mating system shifts and adaptation, because of the availability of a sequenced genome of *C. rubella* (Slotte et al 2013) and because it is possible to generate viable offspring from crosses between *Capsella* species that differ in their mating system (e.g. Slotte et al 2012, Rebernig et al 2015).

In *C. rubella*, the transition to selfing occurred relatively recently (<200 kya), and was associated with speciation from an outcrossing progenitor similar to present-day *C. grandiflora* (Slotte et al. 2013, Foxe et al. 2009, Guo et al. 2009, St Onge et al. 2011, Brandvain et al. 2013). Despite the recent shift to selfing, *C. rubella* already exhibits a derived reduction in petal size and a reduced pollen-ovule ratio, as well as a reduction of the degree of flower opening (Sicard et al 2011, Slotte et al. 2012). *C. rubella* therefore exhibits floral and reproductive characters typical of a selfing syndrome. The selfing syndrome of *C. rubella* is associated with improved efficacy of autonomous self-pollination (Sicard et al. 2011), and regions with quantitative trait loci (QTL) for floral divergence between *C. rubella* and *C. grandiflora* exhibit an excess of fixed differences and reduced polymorphism in *C. rubella* (Slotte et al. 2012). Together, these observations suggest that the rapid evolution of the selfing syndrome in *C. rubella* was driven by positive selection.

While the molecular genetic basis of the selfing syndrome in *C. rubella* has not been identified, it has been suggested that *cis*-regulatory changes could be involved, and a previous study found many flower and pollen development genes to be differentially expressed in flower buds of *C. grandiflora* and *C. rubella* (Slotte et al. 2013). However, these results could be confounded by differences in floral organ sizes and pollen number between *C. rubella* and *C. grandiflora,* and Slotte et al. (2013) did not directly assess *cis*-regulatory changes or investigate possible causes of *cis-*regulatory divergence. There is reason to believe that *cis-*regulatory changes could be partly caused by differences in TE abundance between selfers and outcrossers, as TE silencing can affect nearby gene expression in plants (Hollister and Gaut 2009; Hollister et al. 2011). As *C. rubella* harbors fewer TEs close to genes than *C. grandiflora* (Ågren et al. 2014), this system offers an opportunity to investigate the role of TEs for *cis*-regulatory evolution and for the evolution of floral and reproductive traits in association with the shift to selfing.

In this study we directly assessed *cis*-regulatory divergence by analyzing allele-specific expression in F1 hybrids of *C. grandiflora* and *C. rubella*, and investigated the role of *cis*-regulatory changes for the selfing syndrome in *C. rubella*. We conducted deep sequencing of transcriptomes, small RNAs, as well as genomes of *C. grandiflora* x *C. rubella* hybrids to identify genes with *cis-*regulatory divergence in flower buds and leaves, and tested whether *cis*-regulatory changes in flowers were overrepresented in genomic regions responsible for adaptive phenotypic divergence. We further identifed TEs in *C. rubella* and *C. grandiflora* and tested whether TE insertions targeted by uniquely mapping 24-nt siRNAs were associated with *cis*-regulatory divergence. Our results provide insight into the role of *cis-*regulatory changes in association with the shift to selfing in a wild plant system.

## Results

### Many genes exhibit allele-specific expression in interspecific F1 hybrids

In order to quantify ASE between *C. grandiflora* and *C. rubella*, we generated deep whole transcriptome RNAseq data from flower buds and leaves of three *C. grandiflora* x *C. rubella* F1 hybrids (total 52.1 vs 41.8 Gbp with Q≥30 for flower buds and leaves, respectively). We included three technical replicates for one F1 in order to examine the reliability of our expression data. For all F1s and their *C. rubella* parents, we also generated deep (38-68x) whole genome resequencing data in order to reconstruct parental haplotypes and account for read mapping biases.

F1 RNAseq reads were mapped with high stringency to reconstructed parental haplotypes specific for each F1, i.e. reconstructed reference genomes containing whole-genome haplotypes for both the *C. grandiflora* and the *C. rubella* parent of each F1 (see Material and Methods). We conducted stringent filtering of genomic regions where SNPs were deemed unreliable for ASE analyses due to e.g. high repeat content, copy number variation, or a high proportion of heterozygous genotypes in an inbred *C. rubella* line (for details, see Material and Methods and S1 text); this mainly resulted in removal of pericentromeric regions (S2 Fig - S5 Fig). After filtering, we identified ∼18,200 genes with ∼274,000 transcribed heterozygous SNPs that were amenable to ASE analysis in each F1 (Table 1). The mean allelic ratio of genomic read counts at these SNPs was 0.5 (S6 Fig), suggesting that our bioinformatic procedures efficiently minimized read mapping biases. Furthermore, technical reliability of our RNAseq data was high, as indicated by a mean Spearman’s ρ between replicates of 0.98 (range 0.94-0.99).

**Table 1.** Genes amenable to analysis of ASE in flower bud and leaf samples from the three *C. grandiflora* x *C. rubella* F1s, counts of genes with evidence for ASE and the estimated false discovery rate (FDR) and proportion of genes with ASE.

| F1 designation | Sample | Genes amenable to ASE analysis<sup>a</sup> | Analyzed genes<sup>b</sup> | Heterozygous SNPs in analyzed genes | Genes with ASE PP $\geq 0.95^c$ | FDR | ASE proportion<sup>d</sup> |
| --- | --- | --- | --- | --- | --- | --- | --- |
| Inter3.1 | Flower buds | 18299 | 16857 | 262120 | 4728 | 0.0013 | 0.38 |
| Inter4.1 |  | 18270 | 17837 | 272126 | 5744 | 0.0022 | 0.42 |
| Inter5.1 |  | 18144 | 17448 | 262696 | 5176 | 0.0020 | 0.40 |
| Inter3.1 | Leaves | 18299 | 14877 | 238786 | 5105 | 0.0012 | 0.44 |
| Inter4.1 |  | 18270 | 15784 | 249181 | 8129 | 0.0024 | 0.62 |
| Inter5.1 |  | 18144 | 15478 | 240653 | 4795 | 0.0018 | 0.41 |

<sup>a</sup>Total number of genes with heterozygous SNPs in coding regions remaining after filtering.
<sup>b</sup>Number of genes amenable to ASE analyses with expression data in at least one of the replicates of the sample.
<sup>c</sup>Number of genes with evidence for ASE (posterior probability $\geq 0.95$ ).
<sup>d</sup>Direct estimate of the ASE proportion independent of significance cutoffs.

We assessed ASE using a Bayesian statistical method with a reduced false positive rate compared to the standard binomial test (Skelly et al. 2011). The method uses genomic read counts to model technical variation in ASE and estimates the global proportion of genes with ASE, independent of specific significance cutoffs, and also yields gene-specific estimates of the ASE ratio and the posterior probability of ASE. The model also allows for and estimates the degree of variability in ASE along the gene, through the inclusion of a dispersion parameter.

Based on this method, we estimate that on average, the proportion of assayed genes with ASE is 44.6% (Table 1; S8 Table). In general, most allelic expression biases were moderate, and only 5.9% of assayed genes showed ASE ratios greater than 0.8 or less than 0.2 (Figs. 1 and 2). There was little variation in ASE ratios along genes, as indicated by the distribution of the dispersion parameter estimates having a mode close to zero and a narrow range (Figs. 1 and 2). This suggests that unequal expression of differentially spliced transcripts is not a major contributor to regulatory divergence between *C. rubella* and *C. grandiflora* (Figs. 1 and 2).

**Fig. 1.**
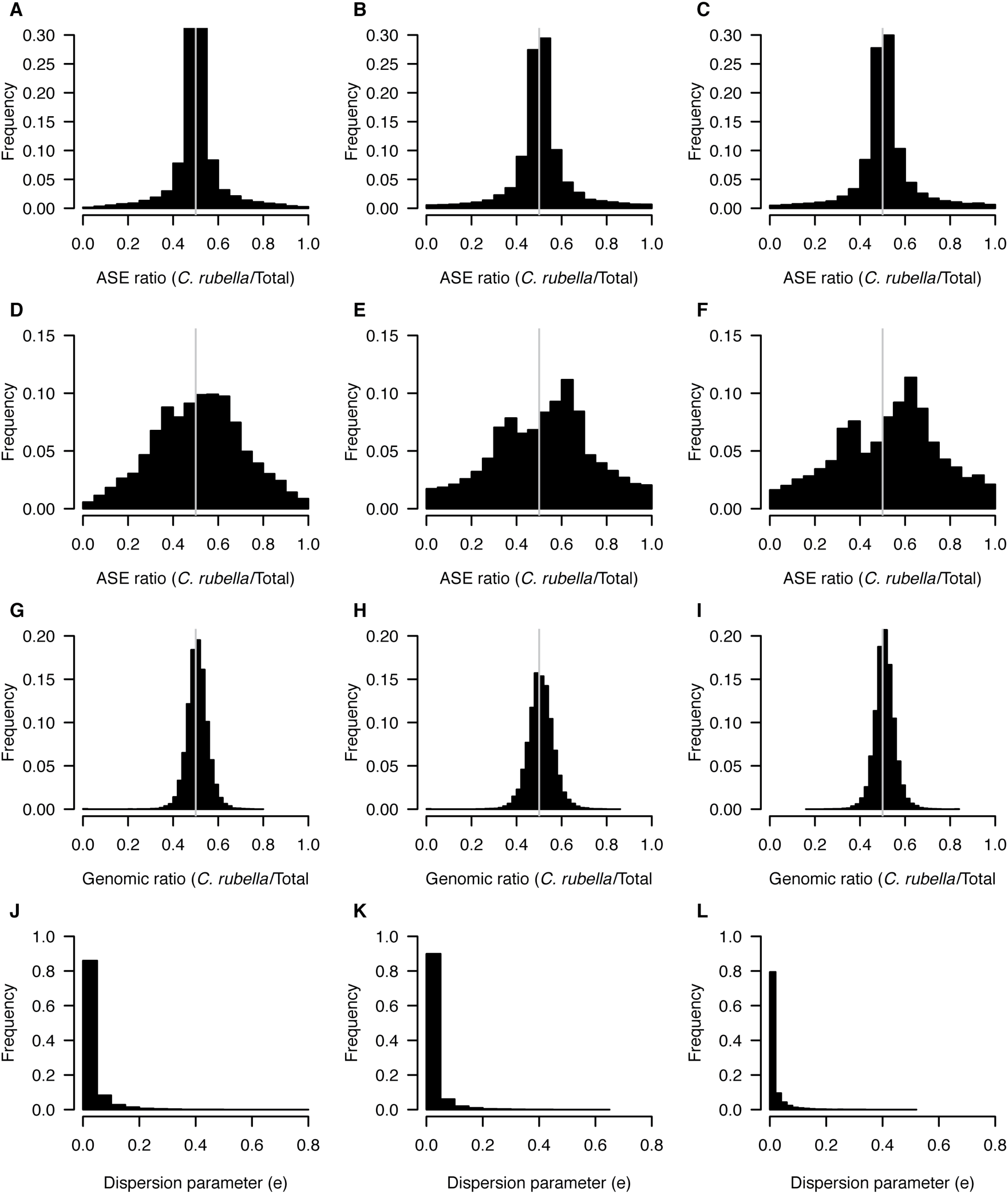
ASE in flower buds. Distributions of ASE ratios (*C. rubella*/Total) for all assayed genes (A, B, C), and for genes with at least 0.95 posterior probability of ASE (D, E, F). Ratio of *C. rubella* to total for genomic reads, for genes with significant ASE (G, H, I), and the distribution of the dispersion parameter that quantifies variability in ASE across genes (J, K, L). All distributions are shown for each of the three interspecific F1s inter 3.1 (left), inter4.1 (middle) and inter5.1 (right).

**Fig. 2.**
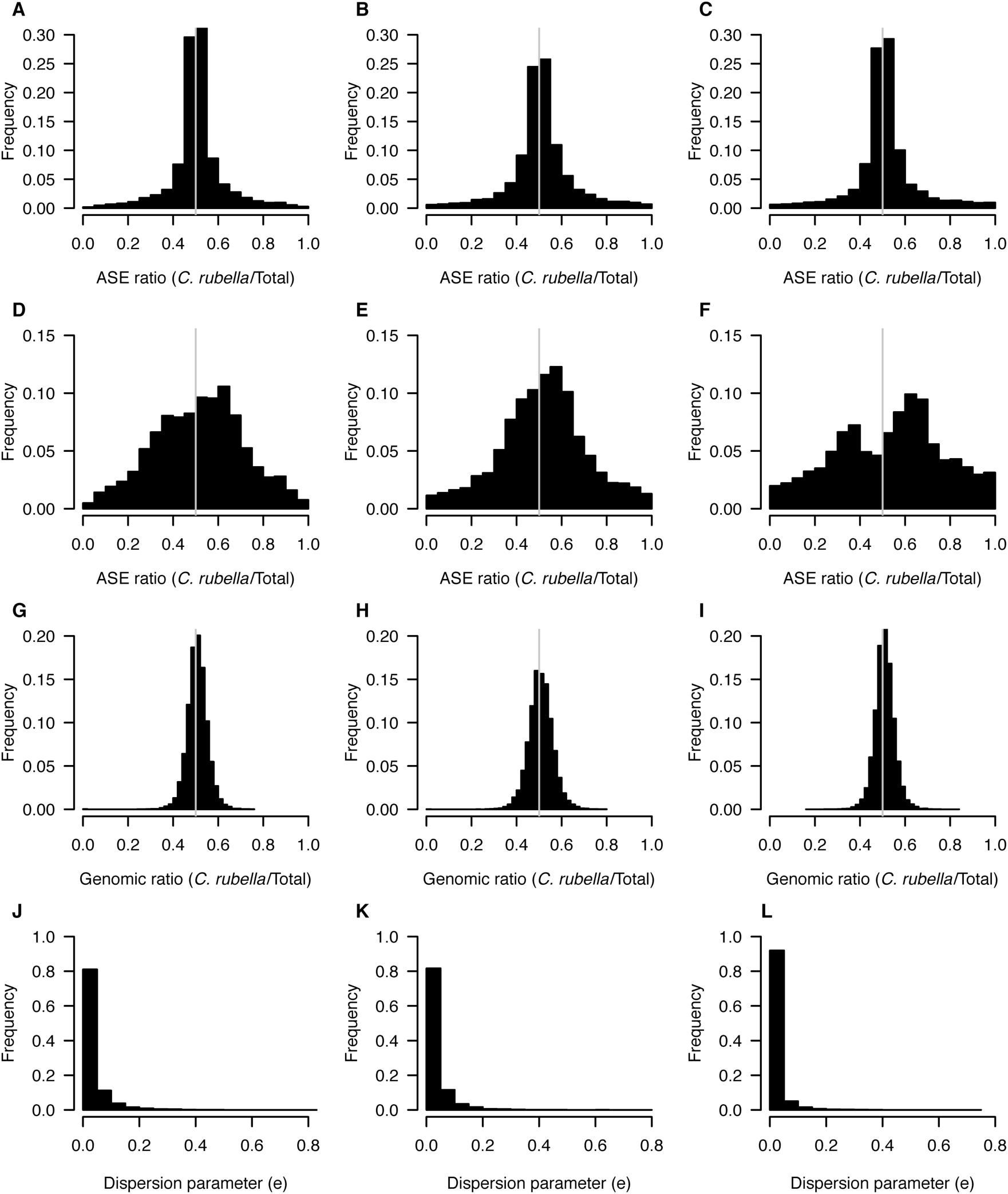
ASE in leaves. Distributions of ASE ratios (*C. rubella*/Total) for all assayed genes (A, B, C), and for genes with at least 0.95 posterior probability of ASE (D, E, F). Ratio of *C. rubella* to total for genomic reads, for genes with significant ASE (G, H, I), and the distribution of the dispersion parameter that quantifies variability in ASE across genes (J, K, L). All distributions are shown for each of the three interspecific F1s inter 3.1 (left), inter4.1 (middle) and inter5.1 (right).

For genes with evidence for ASE (hereafter defined as posterior probability of ASE ≥ 0.95), there was a moderate shift toward higher expression of the *C. rubella* allele (mean ratio *C. rubella*/total=0.56; Figs. 1 and 2). This shift was present for all F1s, for both leaves and flowers (Figs. 1 and 2). No such shift was apparent for genomic reads, and ratios of genomic read counts for SNPs in genes with ASE were very close to 0.5 (mean ratio *C. rubella*/total=0.51; Figs 1 and 2). Furthermore, qPCR with allele-specific probes for five genes validated our ASE results empirically (S9 Table). Thus, *C. rubella* alleles appear to be on average expressed at a higher level than *C. grandiflora* alleles in our F1s.

The mean ASE proportion, as well as the absolute number of genes with ASE was greater for leaves (49%; 6010 genes) than for flower buds (40%; 5216 genes), although this difference was largely driven by leaf samples from one of our F1s (Table 1). Most instances of ASE were specific to either leaves or flower buds, and on average, only 15% of genes expressed in both leaves and flower buds showed consistent ASE in both organs (Fig. 3). Many cases of ASE were also specific to a particular F1, and across all three F1s, there were 1305 genes that showed consistent ASE in flower buds, and 1663 in leaves (Fig. 3).

**Fig. 3.**
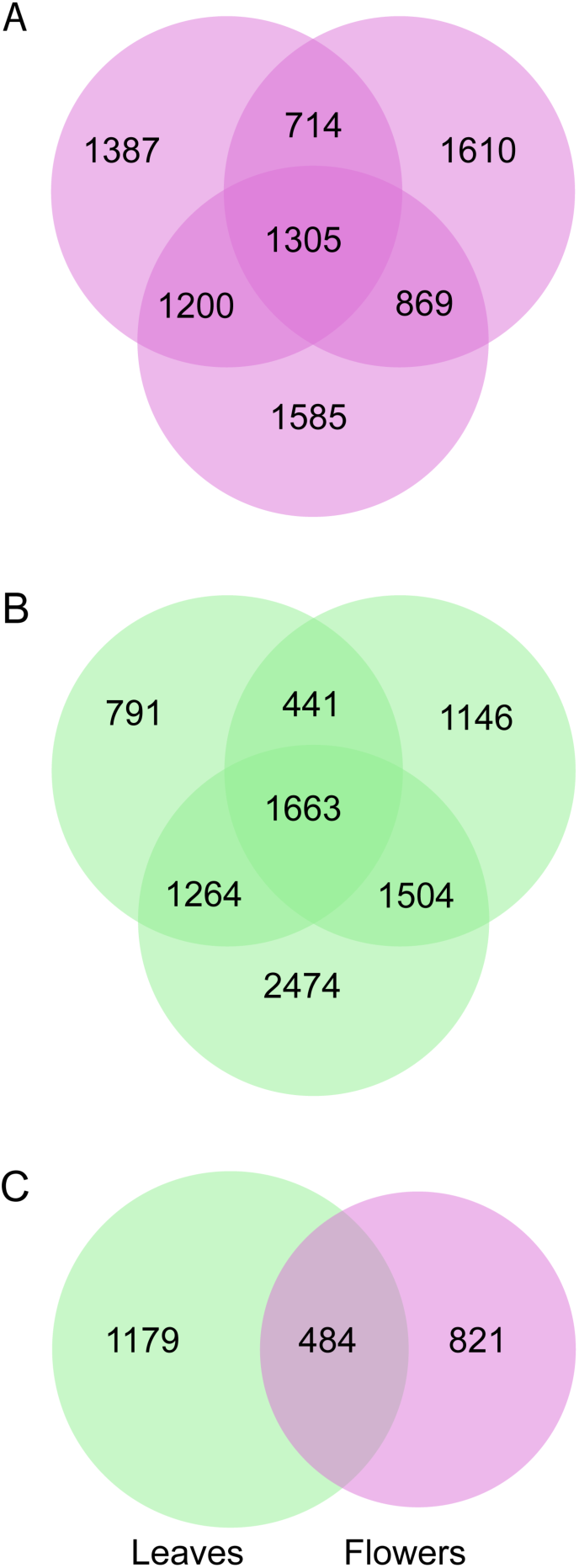
Many cases of ASE are specific to individuals or samples. Venn diagrams showing intersections of genes with ASE in flower buds (A) and leaves (B) of the three F1 individuals, and (C) in all leaf and flower samples, for the set of genes assayed in all F1s.

### Enrichment of *cis-*regulatory changes in genomic regions responsible for phenotypic divergence

We used permutation tests to check for an excess of genes showing ASE within five previously-identified narrow (<2 Mb) QTL regions responsible for floral and reproductive trait divergence (Slotte et al. 2012). These genomic regions harbor major QTL for petal size and flowering time, but also encompass part of the confidence intervals for QTL for sepal size, stamen length and ovule number, as QTL for different floral and reproductive traits are highly overlapping (Slotte et al 2012). As the selfing syndrome has a shared genetic basis in independent *C. rubella* accessions (Sicard et al. 2011, Slotte et al. 2012), we reasoned that genes with consistent ASE across all F1s would be most likely to represent candidate *cis*-regulatory changes underlying QTL. Out of the 1305 genes with ASE in flower buds of all F1s, 85 were found in narrow QTL regions, and this overlap was significantly greater than expected by chance (permutation test, P=0.03; Fig. 4; see Material and Methods for details). In contrast, for leaves, there was no significant excess of genes showing ASE in narrow QTL (permutation test, P=1; Fig. 4). Thus, the association between QTL and ASE in flower buds is unlikely to be an artifact of locally elevated heterozygosity facilitating both ASE and QTL detection, which should affect analyses of both leaf and flower samples.

**Fig. 4.**
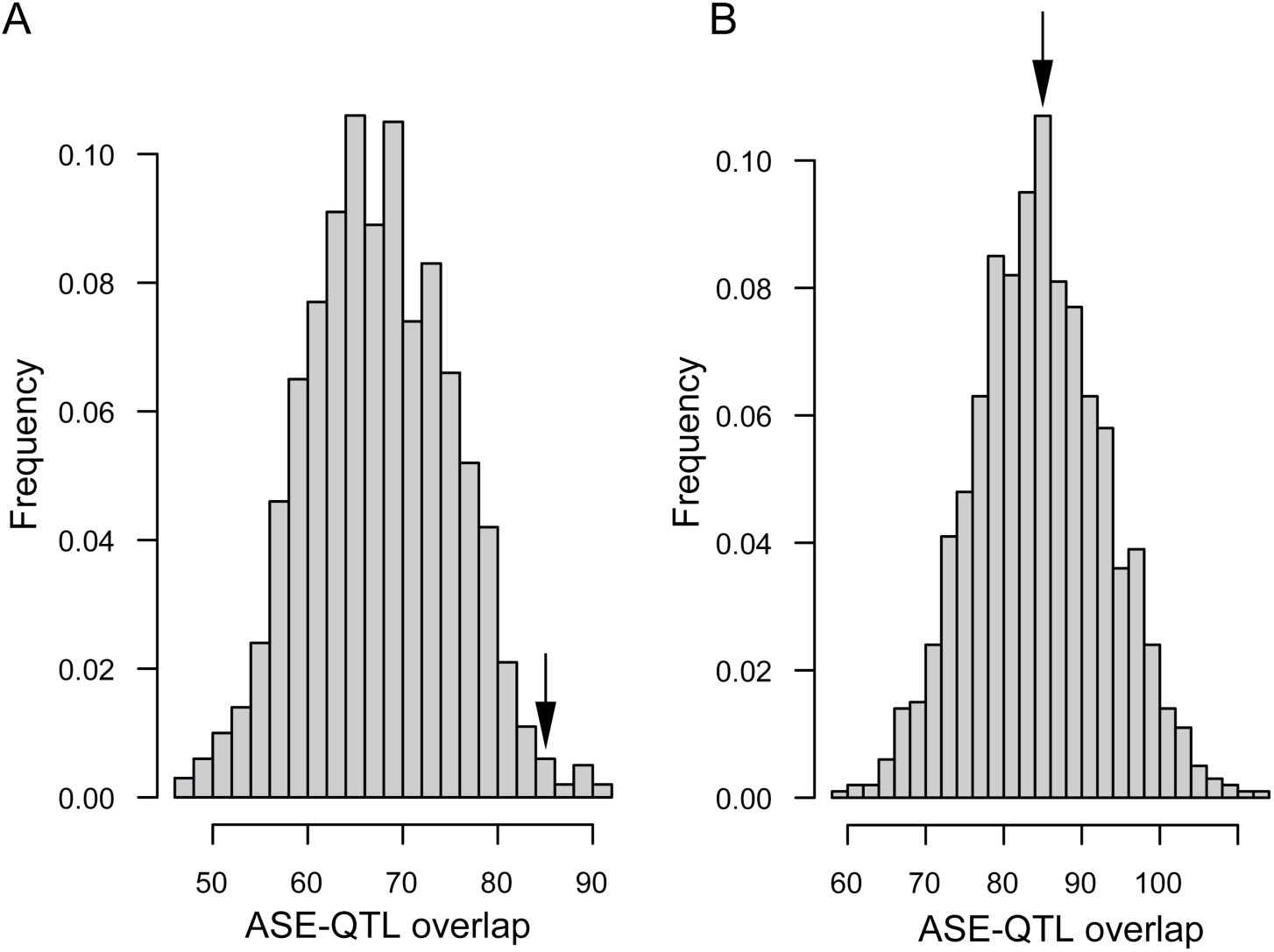
Enrichment of genes with ASE in narrow QTL regions. There is an excess of genes with ASE in narrow QTL regions for flower buds (A) but not for leaves (B). Histograms show the distribution of numbers of genes with ASE that fall within narrow QTL regions, based on 1000 random permutations of the observed number of genes with ASE among all genes where we could assess ASE. Arrows indicate the observed number of genes with ASE that are located in narrow QTL regions.

### List enrichment analyses reveal floral candidate genes with ASE

We conducted list enrichment analyses to characterize the functions of genes showing ASE relative to all genes amenable to analysis of ASE (i.e. harboring heterozygous transcribed SNPs and expressed at detectable levels). There was an enrichment of Gene Ontology (GO) terms involved in defense and stress responses for genes with ASE in flower buds and in leaves (S10 Table). GO terms related to hormonal responses, including brassinosteroid and auxin biosynthetic processes, were specifically enriched among genes with ASE in flower buds (S10 Table). Genes with nearby heterozygous TE insertions were also enriched for a number of GO terms related to reproduction and defense (S11-S12 Table), suggesting that heterozygous TE insertions could be important for patterns of GO term enrichment for genes with ASE

We further identified nineteen genes involved in floral and reproductive development in *A. thaliana*, which are located in QTL regions (see above), and show ASE in flower buds (Table 2). These genes are of special interest as candidate genes for detailed studies of the genetic basis of the selfing syndrome in *C. rubella*.

**Table 2.**
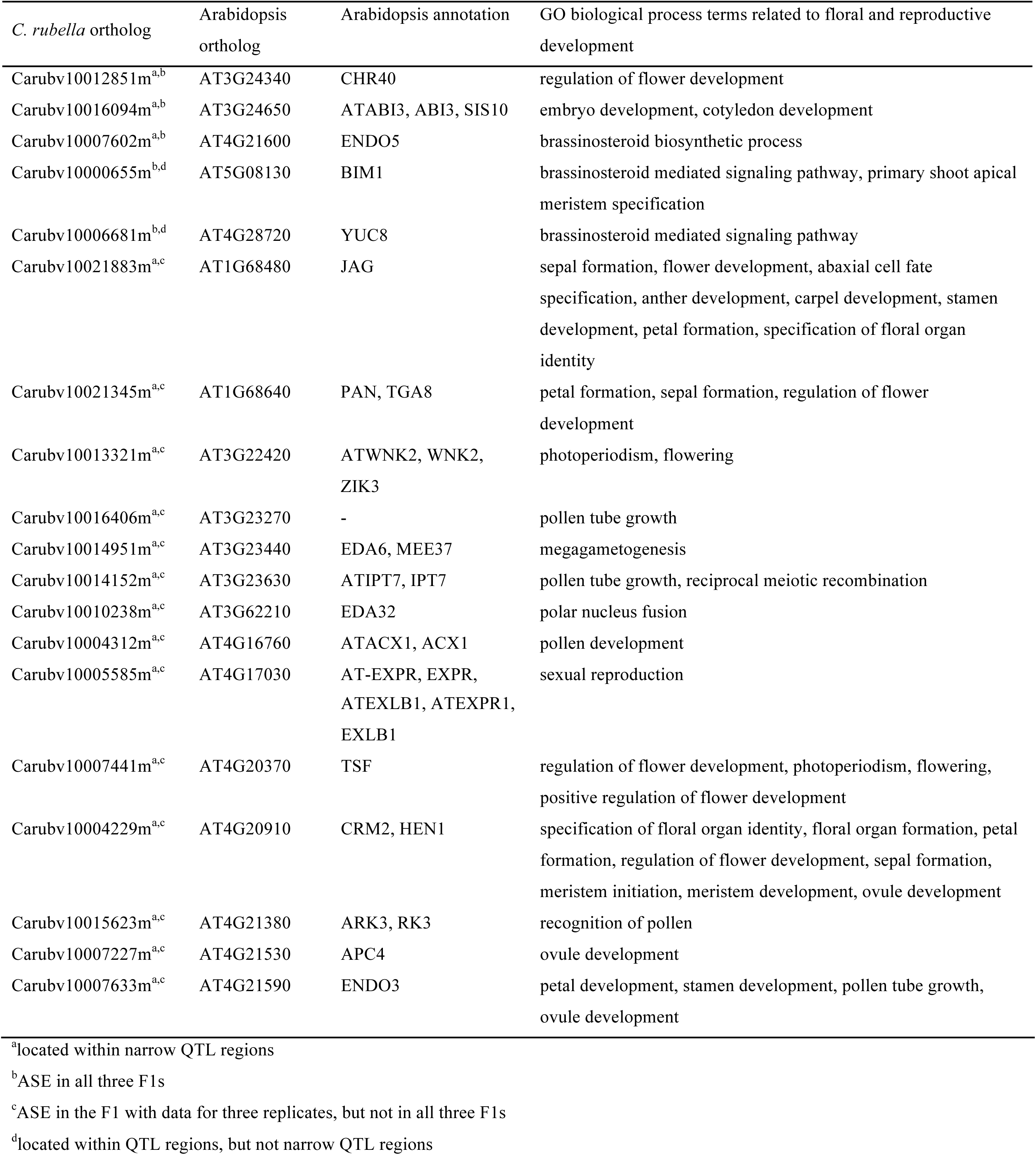
Selfing syndrome candidate genes identified based on ASE, QTL information, and Arabidopsis annotation.

### Intergenic divergence is elevated near genes with ASE

To investigate the role of polymorphisms in regulatory regions for ASE, we assessed levels of heterozygosity in intergenic regions 1 kb upstream of genes, and in previously identified conserved noncoding regions (Williamson et al. 2014) within 5 kb and 10 kb of genes. Genes with ASE were not significantly more likely to be associated with conserved noncoding regions with heterozygous SNPs than genes without ASE. However, levels of intergenic heterozygosity 1 kb upstream of genes were slightly but significantly higher for genes with ASE than for those without ASE (median heterozygosity of 0.016 vs. 0.014, respectively in leaves (Wilcoxon rank sum test, W = 295692325, p-value = 2.26*10^-115^), median heterozygosity of 0.017 vs. 0.014, respectively in flowers (Wilcoxon rank sum test, W = 297625040, p-value = 6.16*10^-142^), S13 Table), suggesting that polymorphisms in regulatory regions upstream of genes might contribute to *cis*-regulatory divergence.

### Enrichment of TEs near genes with ASE

To test whether differences in TE content might contribute to *cis*-regulatory divergence between *C. rubella* and *C. grandiflora*, we examined whether heterozygous TE insertions near genes were associated with ASE. We identified TE insertions specific to the *C. grandiflora* or *C. rubella* parents of our F1s using genomic read data, as in Ågren et al. (2014) (Table 3; see Material and Methods). Overall, we found that *C. rubella* harbored fewer TE insertions close to genes than *C. grandiflora* (on average, 482 vs 1154 insertions within 1 kb of genes in *C. rubella* and *C. grandiflora*, respectively). Among heterozygous TE insertions, *Gypsy* insertions were the most frequent (Table 3); they were also the most frequent genome-wide (Table 3). There was a significant association between heterozygous TE insertions within 1 kb of genes and ASE, for both leaves and flower buds, and the strength of the association was greater for TE insertions closer to genes (Table 4; Fig. 5). This was true for individual F1s, as well as for all F1s collectively (Table 4; Fig. 5; S14 Table).

**Table 3.**
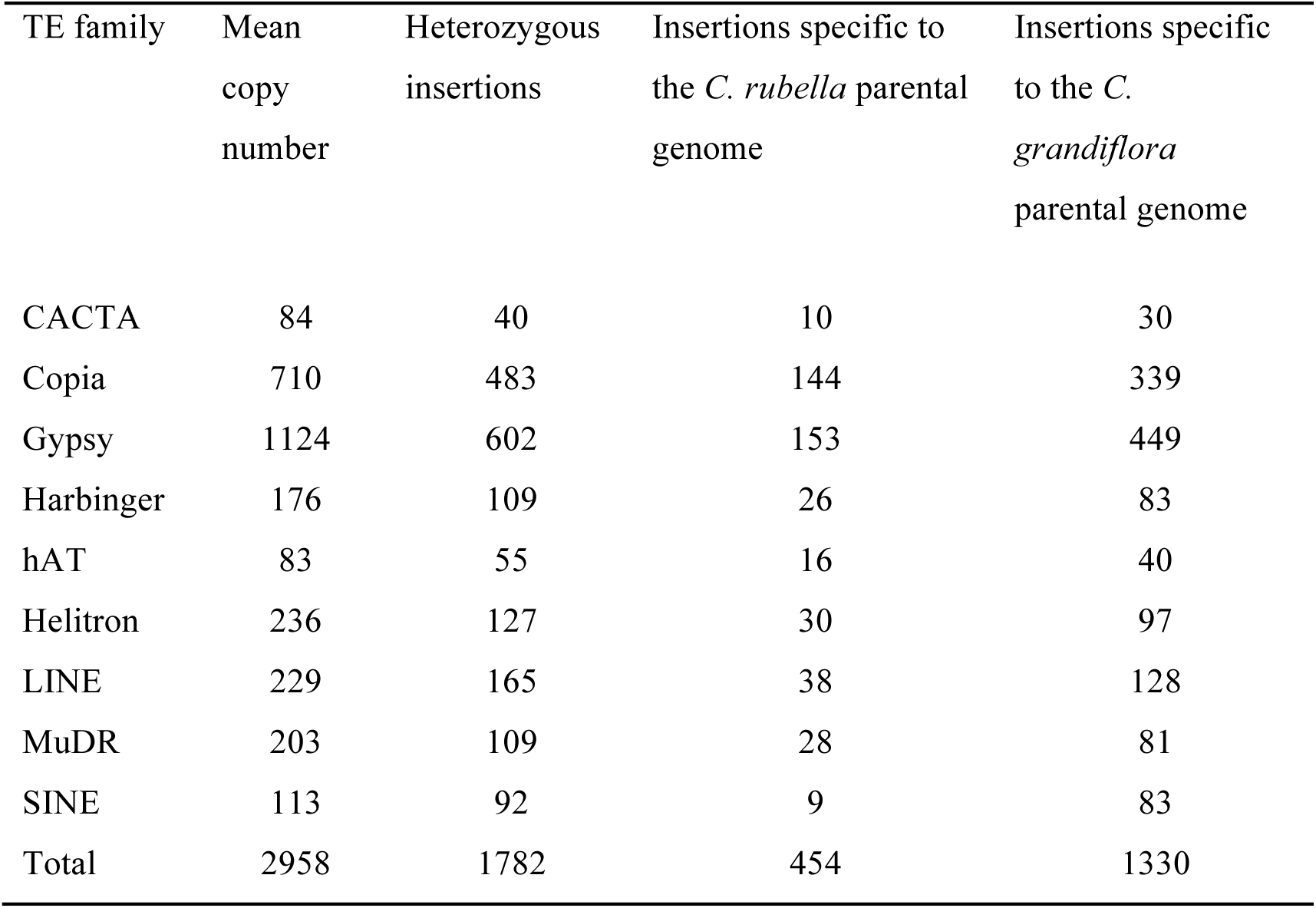
Mean number of TE insertions in three interspecific F1s. The table shows the overall number, as well as heterozygous insertions with parent of origin information.

**Table 4.**
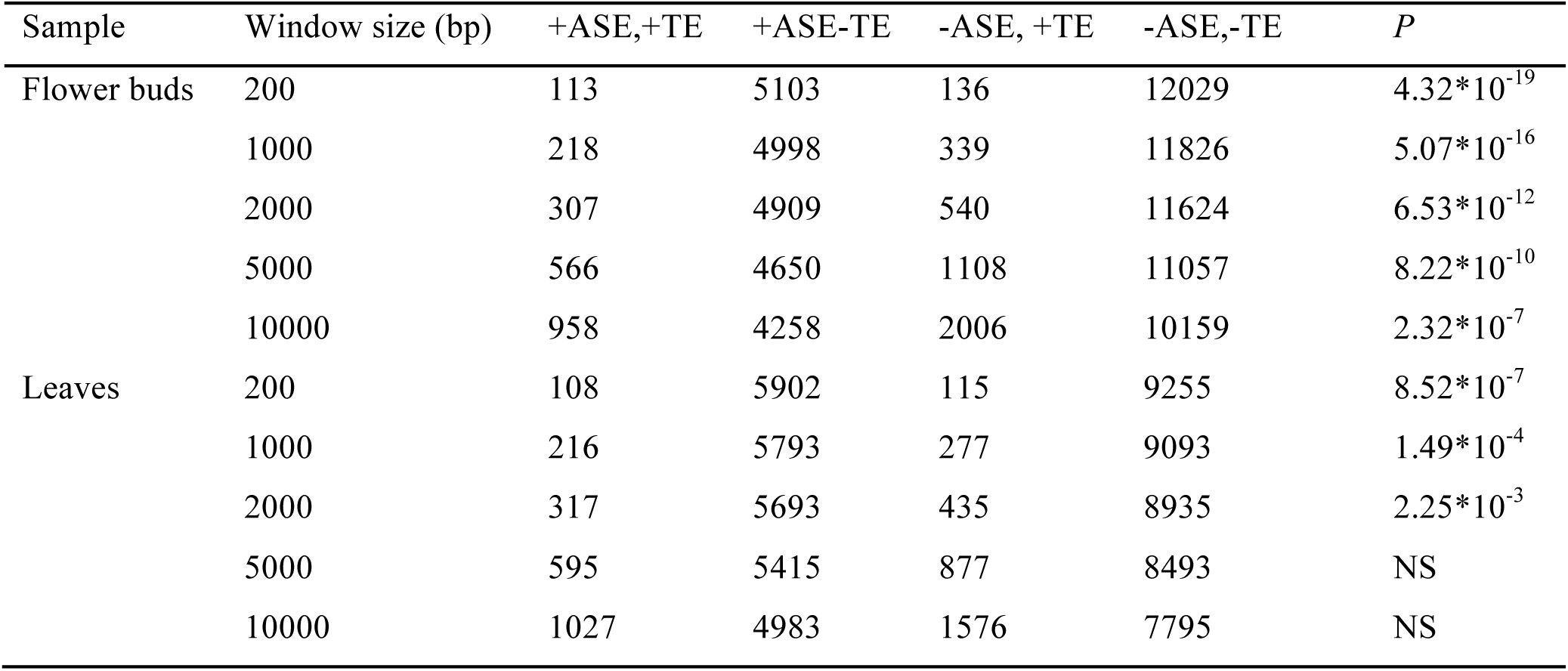
Enrichment of heterozygous TEs near genes with ASE. The table shows mean counts over all three F1s, and Fisher exact test *P*-values. The four categories of counts correspond to numbers of genes with ASE (posterior probability of ASE ≥ 0.95) and TE insertions within a specific window size near the gene (+ASE,+TE), with ASE but without TEs (+ASE,-TE), without ASE but with TE insertions (-ASE,+TE), and with neither ASE nor TEs (-ASE,-TE). NS indicates not significant.

**Fig. 5.**
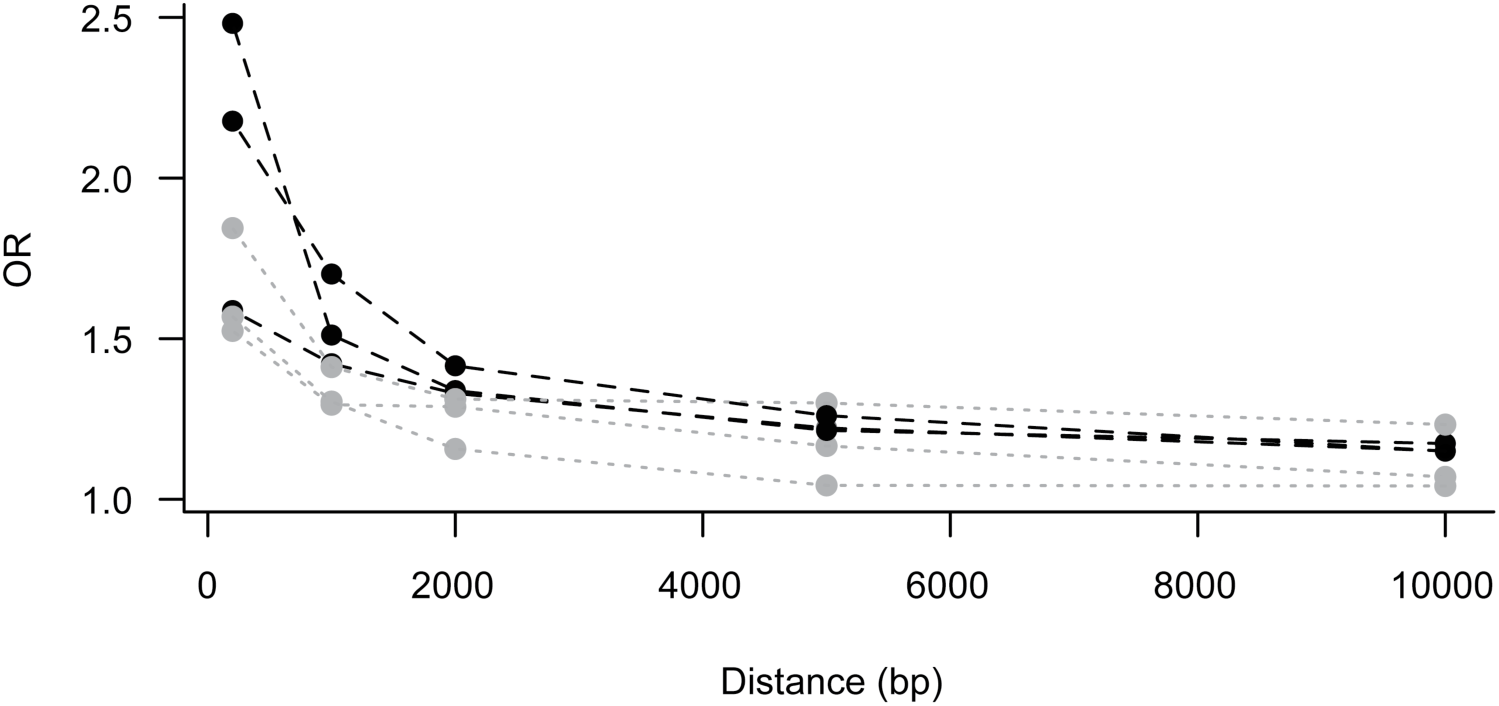
Enrichment of TEs near genes with ASE. The Figure shows odds ratios (OR) of the association between genes with ASE and TEs, with TE insertions scored in four different window sizes (within a distance of 0 bp, 1 kbp, 2 kbp, 5 kbp, and 10 kbp of each gene). Odds ratios for flower buds are shown for all three F1s studies, with values for flower buds in black and leaves in grey.

### TEs targeted by uniquely mapping 24-nt small RNAs are associated with reduced expression of nearby genes

To test whether siRNA-based silencing of TEs might be responsible for the association between TE insertions and ASE in *Capsella*, we analyzed data for flower buds from one of our F1s, for which we had matching small RNA data (see Material and Methods). We selected only those 24-nt siRNA reads that mapped uniquely, without mismatch, to one site within each of our F1s, because uniquely mapping siRNAs have been shown to have a more marked association with gene expression in *Arabidopsis* (Hollister et al. 2009). For each gene, we then assessed the ASE ratio of the allele on the same chromosome as a TE insertion (i.e. ASE ratios were polarized such that relative ASE was equal to the ratio of the expression of the allele with a TE insertion on the same chromosome over the total expression of both alleles), and then further examined the influence of nearby siRNAs.

Overall, the mean relative ASE was reduced for genes with nearby TE insertions (Fig. 6) with a more pronounced effect for TE insertions within 1 kb (within the gene: Wilcoxon rank sum test, W = 1392103, p-value = 8.76*10^-3^; within 200 bp: Wilcoxon rank sum test, W = 1903047, p-value = 7.17*10^-3^; within 1 kb: Wilcoxon rank sum test, W = 3687972, p-value = 8.19*10^-3^). The magnitude of the effect on ASE was more pronounced for genes near TE insertions targeted by uniquely mapping 24-nt siRNAs (Fig. 6; for genes with a TE insertion within the gene: Wilcoxon rank sum test, W = 423369, p-value = 1.36*10^-4^; within 200 bp: W = 540926, p-value = 1.82*10^-5^; within 1 kb: W = 983938, p-value = 3.13*10^-3^). In contrast, no significant effect on ASE was apparent for genes near TE insertions that were not targeted by uniquely mapping 24-nt siRNAs (Fig. 6). Thus, uniquely mapping siRNAs targeting TE insertions appear to be responsible for the association we observe between ASE and TE insertions. Globally, Gypsy and hAT insertions made up a greater proportion of the TE insertions that were targeted by siRNA, compared to those that were not (Chi-squared test, χ=35.9468, P=1.796*10^-5^, Supplementary Figure S7). However, for heterozygous TE insertions within 1 kb of genes there were no significant differences in the composition of TEs that were vs. were not targeted by uniquely mapping siRNAs.

**Fig. 6.**
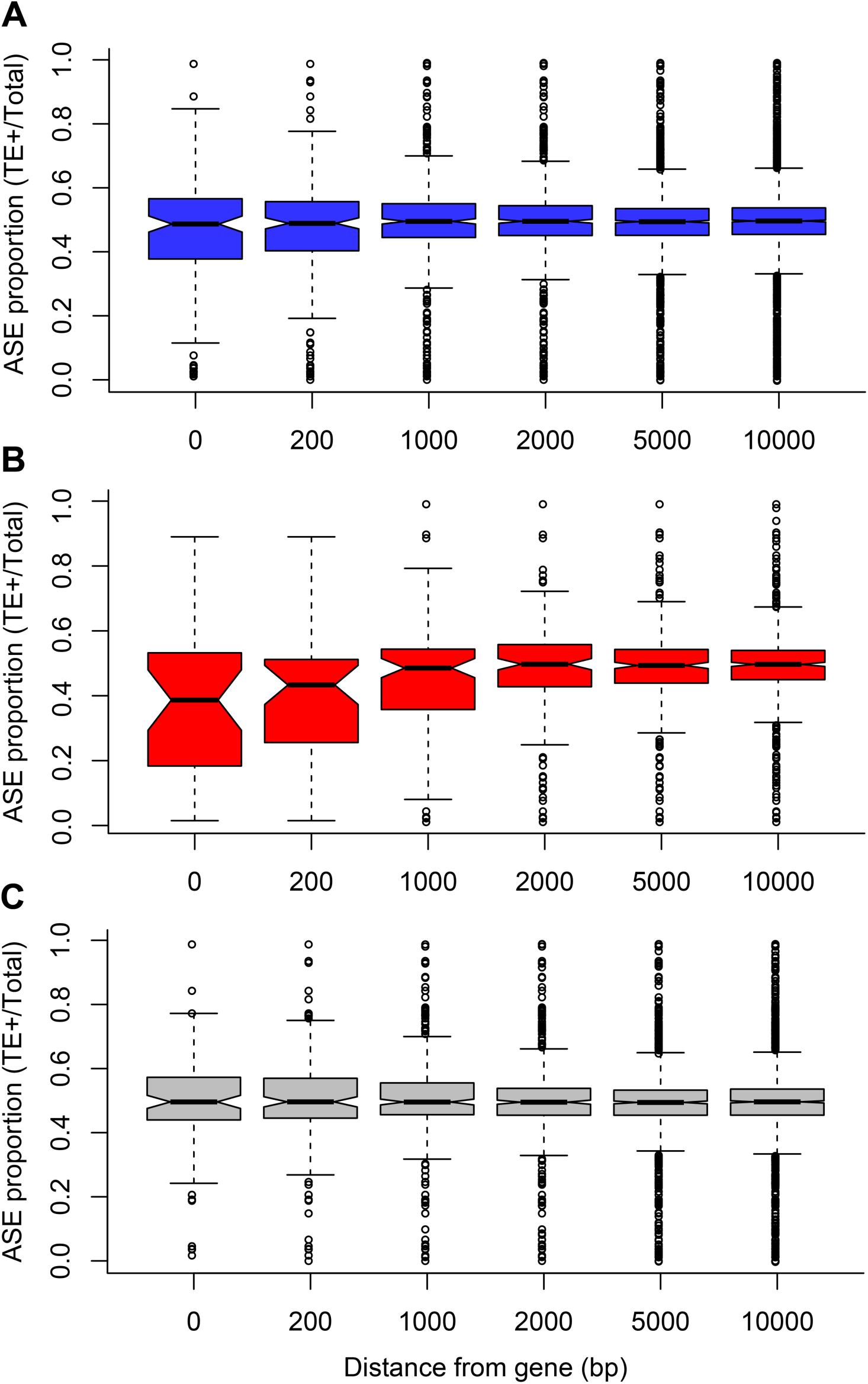
The effect of TE insertions on relative allelic expression. Boxplots show the relative allelic expression (expression of the allele on same haplotype as TE insertion relative to expression of both alleles) for genes near heterozygous TE insertions, scored in a range of window sizes ranging from 0 bp (within the gene) to 10 kbp from the gene. A. The relative allelic expression is reduced for genes with nearby TE insertions. B. The degree of reduction of relative allelic expression is stronger for genes near TE insertions targeted by uniquely mapping siRNA. C. There is no reduction of relative allelic expression for genes near TE insertions that are not targeted by uniquely mapping siRNA.

## Discussion

In this study, we have quantified allele-specific expression in order to understand the role of *cis*-regulatory changes in association with a recent plant mating system shift. Our results indicate that many genes, on average over 40%, harbor *cis*-regulatory differences between *C. rubella* and *C. grandiflora*. The proportion of genes with ASE may seem high given the recent divergence (∼100 kya) between *C. rubella* and *C. grandiflora* (Brandvain et al. 2013, Slotte et al. 2013). However, the majority of genes with ASE showed relatively mild allelic expression biases, and while our estimates are higher than those in a recent microarray-based study of interspecific *Arabidopsis* hybrids (<10%) (He et al. 2012a), our results are consistent with recent analyses of RNAseq data from intraspecific F1 hybrids of *Arabidopsis* accessions (∼30%) (Cubillos et al. 2014). Somewhat higher levels of ASE were found in a recent study of maize and teosinte (∼70% of genes showed ASE in at least one tissue and F1 individual (Lemmon et al. 2014), and using RNAseq data and the same hierarchical Bayesian analysis that we employed, Skelly et al. (2011) estimated that a substantially higher proportion, >70% of assayed genes, showed ASE among two strains of *Saccharomyces cerevisiae.* Thus, our estimates of the proportion of genes with ASE fall within the range commonly observed for recently diverged accessions or lines based on RNAseq data.

Two lines of evidence suggest that *cis-*regulatory changes have contributed to floral and reproductive adaptation to selfing in *C. rubella*. First, we find an excess of genes with ASE in flower buds within previously identified narrow QTL regions for floral and reproductive traits that harbor a signature of selection (Slotte et al. 2012). This suggests either that multiple *cis-*regulatory changes were involved in the evolution of the selfing syndrome in *C. rubella*, or that these regions harbor an excess of *cis-*regulatory changes for other reasons, for instance due to hitchhiking of *cis*-regulatory variants with causal variants for the selfing syndrome. Distinguishing between these hypotheses will require identification of causal genetic changes for the selfing syndrome in *C. rubella*. In contrast, no such excess is present for genes with ASE in leaves, suggesting that this observation is not simply a product of higher levels of divergence among *C. rubella* and *C. grandiflora* in certain genomic regions facilitating both QTL delimitation and ASE analysis. Second, we find that genes involved in hormonal responses, including brassinosteroid biosynthesis, are overrepresented among genes with ASE in flower buds, but not in leaves. Based on a study of differential expression and functional information from *Arabidopsis thaliana*, regulatory changes in this pathway were previously suggested to be important for the selfing syndrome in *C. rubella* (Slotte et al. 2013). While we do not identify ASE at the same genes as in Slotte et al. 2013, our work nonetheless provides support for *cis-*regulatory changes at other genes in the brassinosteroid pathway contributing to the selfing syndrome of *C. rubella*. Future studies should conduct fine-scale mapping and functional validation to fully explore this hypothesis. To facilitate this work, we have identified a set of candidate genes with ASE that are located in genomic regions harboring QTL for floral and reproductive trait divergence between *C. rubella* and *C. grandiflora*. Of particular interest in this list is the gene *JAGGED* (*JAG*), which is involved in determining petal growth and shape by promoting cell proliferation in *A. thaliana* (Sauret-Güeto et al. 2013, Schiessl et al. 2014). As *C. rubella* has reduced petal size due to a shortened period of proliferative growth (Sicard et al. 2011), and the *C. rubella* allele is expressed at a lower level than the *C. grandiflora* allele, this gene is a very promising candidate gene for the selfing syndrome.

Our work also provides general insights into the nature of *cis-*regulatory divergence. Indeed, many instances of ASE were specific to a particular individual or tissue, an observation also supported by recent studies (e.g. Lemmon et al. 2014, He et al. 2012a). This suggests that there is substantial variation in ASE depending on genotype and developmental stage, consistent with the reasoning that *cis*-regulatory changes can have very specific effects, but expression noise is probably also a contributing factor. It is also difficult to completely rule out the possibility that some cases of subtle ASE may not represent biologically meaningful *cis*-regulatory variation. However, in our analyses, we took several steps to model and account for technical variation in order to reduce the incidence of false positives. We also cannot fully rule out imprinting effects as potential causes of ASE, because generating reciprocal F1 hybrids was not possible due to seed abortion in *C. rubella* x *C. grandiflora* crosses. However, we do not expect these effects to make a major contribution to the patterns we observed; in *Arabidopsis*, imprinting effects are only prevalent in endosperm tissue, and are rare in more advanced stage tissues such as those analyzed here (Scott et al. 1998, Wolff et al. 2011, Cubillos et al. 2014), which suggests that imprinting is not likely to be responsible for the patterns we observe.

One somewhat unexpected finding was the global shift in expression levels toward higher relative expression of the *C. rubella* allele in the F1 hybrids. No marked bias was present for the same SNPs and genes in our genomic data, suggesting that if systematic bioinformatic biases are the cause, the effect is specific to transcriptomic reads. This seems unlikely to completely explain the shift in expression that we observe, as we made considerable effort to avoid reference mapping bias, including high stringency mapping of transcriptomic reads to reconstructed parental haplotypes specific to each F1. Similar global shifts toward higher expression of the alleles from one parent have also been observed in F1s of maize and teosinte (Lemmon et al. 2014) and *Drosophila* (McManus et al. 2010). An even stronger bias toward higher expression of the *A. lyrata* allele was recently observed in F1s of *A. thaliana* and *A. lyrata* (He et al. 2012a), and was attributed to interspecific differences in gene silencing. Our results mirror those seen in some allopolyploids, where homeologs from one parental species can be expressed at a markedly higher level than those from the other parental species (e.g. Chang et al 2010; Flagel & Wendel 2010; Schnable et al 2011; Yoo et al. 2013).

To investigate potential mechanisms for *cis*-regulatory divergence, we first examined heterozygosity in regulatory regions and conserved noncoding regions close to genes. While genes with ASE in general showed slightly elevated levels of heterozygosity upstream of genes, there was no enrichment of conserved noncoding regions with heterozygous SNPs close to genes with ASE. It thus seems likely that divergence in regulatory regions in the proximity of genes, but not specifically in conserved noncoding regions, has contributed to global *cis*-regulatory divergence between *C. rubella* and *C. grandiflora*.

To examine biological explanations for the shift toward a higher relative expression of *C. rubella* alleles, we examined the relationship between TE insertions and ASE. As *C. rubella* harbors a lower number of TE insertions near genes than *C. grandiflora,* we reasoned that TE silencing might contribute to the global shift in expression toward higher relative expression of the *C. rubella* allele, with *C. grandiflora* alleles being preferentially silenced due to targeted methylation of nearby TEs, through transcriptional gene silencing mediated by 24-nt siRNAs. Our results are consistent with this hypothesis. Not only is there is an association between genes with TEs and heterozygous TE insertions in our F1s, there is also reduced expression of alleles that reside on the same haplotype as a nearby TE insertion, and the reduction is particularly strong for TEs that are targeted by uniquely mapping siRNAs. In contrast, no effect on ASE is apparent for TEs that are not targeted by uniquely mapping siRNAs. Moreover, the relatively limited spatial scale over which siRNA-targeted TE insertions are associated with reduced expression of nearby genes (<1 kb) is consistent with previous results from *Arabidopsis* (Hollister et al. 2009, Hollister et al. 2011, Wang et al. 2013). Our findings therefore suggest that silencing of TE insertions close to genes is important for global *cis*-regulatory divergence between *C. rubella* and *C. grandiflora*.

Why then do *C. rubella* and *C. grandiflora* differ with respect to silenced TEs near genes? In *Arabidopsis*, methylated TE insertions near genes appear to be predominantly deleterious, and exhibit a signature of purifying selection (Hollister et al. 2009). The reduced prevalence of TE insertions near genes in *C. rubella* could be caused by rapid purging of recessive deleterious alleles due to increased homozygosity as a result of self-fertilization (Arunkumar et al. 2014). However, we prefer the alternative interpretation that deleterious alleles that were rare in the outcrossing ancestor were preferentially lost in *C. rubella*, mainly as a consequence of the reduced effective population size associated with the shift to selfing. This is in line with analyses of polymorphism and divergence at nonsynonymous sites, for which *C. rubella* exhibits patterns consistent with a general relaxation of purifying selection (Slotte et al. 2013).

If TE dynamics are generally important for *cis*-regulatory divergence in association with plant mating system shifts, we might expect different effects on *cis-*regulatory divergence depending not only on the genome-wide distribution of TEs, but also on the efficacy of silencing mechanisms in the host (Hollister et al. 2009, Hollister et al. 2011, Ågren et al. 2015). For instance, He et al. (2012a) found a shift toward higher relative expression of alleles from the outcrosser *A. lyrata,* which harbors a higher TE content, a fact which they attributed to differences in silencing efficacy between *A. thaliana* and *A. lyrata*; indeed, TEs also showed upregulation of the *A. lyrata* allele (He et al. 2012b) and *A. lyrata* TEs were targeted by a lower fraction of uniquely mapping siRNAs (Hollister et al. 2011). In contrast, we found no evidence for a difference in silencing efficacy between *C. rubella* and *C. grandiflora*, which harbor similar fractions of uniquely mapping siRNAs (12% vs 10% uniquely mapping/total 24-nt RNA reads for *C. rubella* and *C. grandiflora*, respectively). Thus, in the absence of strong divergence in silencing efficacy, differences in the spatial distribution of TEs, such as those we observe between *C. rubella* and *C. grandiflora*, might be more important for *cis*-regulatory divergence. More studies of ASE in F1s of selfers of different ages and their outcrossing relatives are needed to assess the general contribution of differences in silencing efficacy versus genomic distribution of TE insertions for *cis*-regulatory divergence in association with mating system shifts.

## Conclusions

We have shown that many genes exhibit *cis*-regulatory changes between *C. rubella* and *C. grandiflora* and that there is an enrichment of genes with floral ASE in genomic regions responsible for phenotypic divergence. In combination with analyses of the function of genes with floral ASE, this suggests that *cis-*regulatory changes have contributed to the evolution of the selfing syndrome in *C. rubella*. We further observe a general shift toward higher relative expression of the *C. rubella* allele, an observation that can in part be explained by elevated TE content close to genes in *C. grandiflora* and reduced expression of *C. grandiflora* alleles due to silencing of nearby TEs. These results support the idea that TE dynamics and silencing are of general importance for *cis*-regulatory divergence in association with plant mating system shifts.

## Material and Methods

### Plant material

We generated three interspecific *C. grandiflora* x *C. rubella* F1s by crossing two accessions of the selfer *C. rubella* as pollen donor with three accessions of the outcrosser *C. grandiflora* as seed parent (S16 Table). No viable seeds were obtained from reciprocal crosses. Seeds from F1s and their *C. rubella* parental lines were surface-sterilized and germinated on 0.5 x Murashige-Skoog medium. We transferred one-week old seedlings to soil in pots that were placed in randomized order in a growth chamber (16 h light: 8 h dark; 20° C: 14° C). After four weeks, but prior to bolting, we sampled young leaves for RNA sequencing. Mixed-stage flower buds were sampled 3 weeks later, when all F1s were flowering. To assess data reliability, we collected three separate samples of leaves and flower buds from one F1 individual, and three biological replicates of one *C. rubella* parental line. For genomic DNA extraction, we sampled leaves from all three F1 individuals as well as from their *C. rubella* parents. For small RNA sequencing, we germinated six F2 offspring from one of our F1 individuals and sampled flower buds as described above.

### Sample preparation and sequencing

We extracted total RNA for whole transcriptome sequencing with the RNEasy Plant Mini Kit (Qiagen, Hilden, Germany). For small RNA sequencing, we extracted total RNA using the mirVana kit (Life Technologies). For whole genome sequencing, we used a modified CTAB DNA extraction (Doyle and Doyle 1987) to obtain predominantly nuclear DNA. RNA sequencing libraries were prepared using the TruSeq RNA v2 protocol (Illumina, San Diego, CA, USA). DNA sequencing libraries were prepared using the TruSeq DNA v2 protocol. Small RNA libraries were prepared from 1 µg of total RNA using the TruSeq SmallRNA SamplePrep fom Illumina according to the manufacturer’s protocol (#15004197 rev E; Illumina, San Diego, CA, USA). Sequencing was performed on an Illumina HiSeq 2000 instrument (Illumina, San Diego, CA, USA) to gain 100bp paired end reads, except for small RNA samples for which single end 50 bp reads were obtained. Sequencing was done at the Uppsala SNP & SEQ Technology Platform, Uppsala University, except for accession *C. rubella* Cr39.1 where genomic DNA sequencing was done at the Max Planck Institute of Developmental Biology, Tübingen. In total, we obtained 93.9 Gbp (Q≥30) of RNAseq data, with an average of 9.3 Gbp per sample. In addition we obtained 45.6 Gbp (Q≥30) of DNAseq data, corresponding to a mean expected coverage per individual of 52x, and 106,110,000 high-quality (Q≥30) 50 bp small RNA reads. All sequence data has been submitted to the European Bioinformatics Institute (www.ebi.ac.uk), with study accession number: PRJEB9020.

### Sequence quality and trimming

We merged read pairs from fragment spanning less than 185 nt (this also removes potential adapter sequences) in SeqPrep (https://github.com/jstjohn/SeqPrep) and trimmed reads based on sequence quality (phred cutoff of 30) in CutAdapt 1.3 (Martin 2011). For DNA and RNAseq reads, we removed all read pairs where either of the reads was shorter than 50 nt. We then analyzed each sample individually using fastQC v. 0.10.1 (http://www.bioinformatics.babraham.ac.uk/projects/fastqc/) to identify potential errors that could have occurred in the process of amplifying DNA and RNA. We assessed RNA integrity by analyzing the overall depth of coverage over annotated coding genes, using geneBody_coverage.py that is part of the RSeQC package v. 2.3.3 (Wang et al. 2012). For DNA reads we analyzed the genome coverage using bedtools v.2.17.0 (Quinlan and Hall 2010) and removed all potential PCR duplicates using Picard v.1.92 (http://picard.sourceforge.net). Small RNA reads were trimmed using custom scripts and CutAdapt 1.3 and filtered to retain only reads of 24 nt length.

### Read mapping and variant calling

We mapped both genomic reads and RNAseq reads to the v1.0 reference *C. rubella* assembly (Slotte et al. 2013) (http://www.phytozome.net/capsella). For RNAseq reads we used STAR v.2.3.0.1 (Dobin et al. 2013) with default parameters. For genomic reads we modified the default STAR settings to avoid splitting up reads, and for mapping 24-nt small RNA we used STAR with settings modified to require perfect matches to the parental haplotypes of the F1s as well as to a TE library based on multiple Brassicaceae species and previously used in Slotte et al. (2013).

Variant calling was done in GATK v. 2.5-2 (McKenna et al. 2010) according to GATK best practices (DePristo et al. 2011, Van der Auwera et al. 2013). Briefly, after duplicate marking, local realignment around indels was undertaken, and base quality scores were recalibrated, using a set of 1,538,085 SNPs identified in *C. grandiflora* (Williamson et al. 2014) as known variants. Only SNPs considered high quality by GATK were kept for further analysis. Variant discovery was done jointly on all samples using the UnifiedGenotyper, and for each F1, genotypes were phased by transmission, by reference to the genotype of its highly inbred *C. rubella* parental accession.

We validated our procedure for calling variants in genomic data by comparing our calls for the inbred line *C. rubella* 1GR1 at 176,670 sites sequenced in a different individual from the same line by Sanger sequencing (Slotte et al. 2010). Overall, we found 29 calls that differed among the two sets, resulting in an error rate of 0.00016, considerably lower than the level of divergence among *C. rubella* and *C. grandiflora* (0.02; Brandvain et al. 2013).

### Reconstruction of parental haplotypes of interspecific F1s

We reconstructed genome-wide parental haplotype sequences for each interspecific F1 and used these as a reference sequence for mapping genomic and transcriptomic reads for ASE analyses. This was done to reduce effects of read mapping biases on our analyses of ASE by increasing the number of mapped reads and reducing mismapping that can result when masking heterozygous SNPs in F1s (Degner et al. 2009).

To reconstruct parental genomes for each F1, we first conducted genomic read mapping, variant calling and phasing by reference to the inbred *C. rubella* parent as described in the section “Read Mapping and Variant Calling” above. The resulting phased vcf files were used in conjunction with the *C. rubella* reference genome sequence to create a new reference for each F1, containing both of its parental genome-wide haplotypes. Read mapping of both genomic and RNA reads from each F1 was then redone to its specific parental haplotype reference genome, and read counts at all reliable SNPs (see section “Filtering” below) were obtained using Samtools mpileup and a custom software written in javascript by Johan Reimegård. The resulting files with allele counts for genomic and transcriptomic data were used in all downstream analyses of allelic expression biases (see section “Analysis of Allele-Specific Expression” below).

### Filtering

We used two approaches to filter the genome assembly to identify regions where we have high confidence in our SNP calls. Genomic regions with evidence for large-scale copy number variation were identified using Control-FREEC (Boeva et al. 2011), and repeats and selfish genetic elements were identified using RepeatMasker 4.0.1 (http://www.repeatmasker.org). Additionally, we identified genomic regions with unusually high proportions of heterozygous genotype calls in a lab-inbred *C. rubella* line, which is expected to be highly homozygous. Regions with evidence for high proportions of repeats, copy number variation or high proportion of heterozygous calls in the inbred line mainly corresponded to centromeric and pericentromeric regions, and these were removed from consideration in further analyses of allele-specific expression (S2 Fig. - S5 Fig.).

### Analysis of allele-specific expression

Analyses of allele-specific expression (ASE) were done using a hierarchical Bayesian method developed by Skelly et al. (2011). The method requires read counts at heterozygous coding SNPs for both genomic and transcriptomic data. Genomic read counts are used to fit the parameters of a beta-binomial distribution, in order to obtain an empirical estimate of the distribution of variation in allelic ratios due to technical variation (as there is no true ASE for genomic data on read counts for heterozygous SNPs). This distribution is then used in analyses of RNAseq data where genes are assigned posterior probabilities of exhibiting ASE.

We conducted ASE analyses using the method of Skelly et al. (2011) for each of our three F1 individuals. Prior to analyses, we filtered the genomic data to only retain read counts for heterozygous SNPs in coding regions that did not overlap with neighboring genes, and following Skelly et al. (2011), we also removed SNPs that were the most strongly biased in the genomic data (specifically, in the 1% tails of a beta-binomial distribution fit to all heterozygous SNPs in each sample), as such highly biased SNPs may result in false inference of variable ASE if retained. The resulting data set showed very little evidence for read mapping bias affecting allelic ratios: the mean ratio of *C. rubella* alleles to total was 0.507 (S6 Fig).

All analyses were run in triplicate and MCMC convergence was checked by comparing parameter estimates across independent runs from different starting points, and by assessing the degree of mixing of chains. For all analyses of RNA counts, we used median estimates of the parameters of the beta-binomial distribution from analyses of genomic data for all three F1s (S8 Table). Runs were completed on a high-performance computing cluster at Uppsala University (UPPMAX) using the pqR implementation of R (http://www.pqr-project.org), for 200,000 generations or a maximum runtime of 10 days. We discarded the first 10% of each run as burn-in prior to obtaining parameter estimates.

### ASE validation by qPCR

We validated ASE results by performing qPCR with TaqMan® Reverse Transcription Reagents (LifeTechnologies, Carlsbad, CA, USA) using oligo(dT)_16_s to convert mRNA into cDNA using the manufacturers protocol and performed qPCR with the Custom TaqMan® Gene Expression Assay (LifeTechnologies, Carlsbad, CA, USA) with the colors FAM and VIC using manufacturers protocol. The qPCR for both alleles was multiplexed in one well to directly compare the two alleles using a Bio-Rad CFX96 Touch™ Real-Time PCR Detection System (Bio-Rad, Hercules, CA, USA). To exclude color bias, we used reciprocal probes with VIC and FAM colorant (S15 Table). The expression difference between the *C. rubella* and *C. grandiflora* allele was quantified using the difference in relative expression between the two alleles, as well as the Quantification Cycle (Cq value). A lower Cq value correlates with a higher amount of starting material in the sample. If the direction of allelic imbalance inferred by qPCR was the same as for ASE inferred by the method by Skelly et al. (2011), we considered that the qPCR supported the ASE results. For further details see S1 Text.

### Enrichment of genes with ASE in genomic regions responsible for phenotypic divergence

We tested whether there was an excess of genes with evidence for ASE (posterior probability of ASE ≥ 0.95 in all three F1 hybrids) in previously identified genomic regions harboring QTL for phenotypic divergence between *C. rubella* and *C. grandiflora* (Slotte et al. 2012). For this purpose, we concentrated on narrow QTL regions, defined as in a previous study (Slotte et al. 2012) (i.e. QTL regions with 1.5-LOD confidence intervals <2 Mb). The five QTL regions that met our criteria for inclusion as narrow QTL were non-overlapping and corresponded to previously identified QTL for floral and reproductive traits (on scaffolds 2 and 7 for petal width, on scaffold 7 for petal length and on scaffolds 1 and 3 for flowering time). As QTL for floral and reproductive traits are generally highly overlapping these genomic regions also encompass part of the confidence intervals for other QTL, including a major QTL for petal length on scaffold 2, and QTL for sepal length, stamen length and ovule number on scaffold 7). Significance was based on a permutation test (1000 permutations) in R 3.1.2.

### List enrichment tests of GO terms

We tested for enrichment of GO biological process terms using Fisher exact tests in the R package TopGO (Alexa et al. 2006). GO terms were downloaded from TAIR (http://www.arabidopsis.org) on September 3rd, 2013, for all *A. thaliana* genes that have orthologs in the *C. rubella* v1.0 annotation, and we only considered GO terms with at least two annotated members in the background set.

We tested for enrichment of GO biological process terms among genes with ASE in all of our F1s Separate tests were conducted for leaf and flower bud samples, and background sets consisted of all genes where we could assess ASE in either leaves or flower buds.

We used the same approach to test for enrichment of GO biological process terms among genes within 1 kb and 2 kb of heterozygous TE insertions in F1 Inter4.1, for which we had matching small RNA data. For this purpose, separate tests were done for all heterozygous TE insertions, heterozygous TE insertions targeted by uniquely mapping siRNAs, and heterozygous TE insertions not targeted by siRNAs. For these tests, the background sets consisted of all annotated *C. rubella* genes.

### Intergenic heterozygosity in regulatory and conserved noncoding regions

We quantified intergenic heterozygosity 1 kb upstream of genes using VCFTools (Danecek et al. 2011), and compared levels of polymorphism among genes with and without ASE using a Wilcoxon rank sum test. We further assessed whether there was an enrichment of conserved noncoding elements (identified in Williamson et al. (2014)) with heterozygous SNPs within 5 kb of genes with ASE, using Fisher exact tests. Separate tests were conducted for each F1.

### Identification of TE insertions and association with ASE

We used PoPoolationTE (Kofler et al. 2012) to identify transposable elements in our F1s. While intended for pooled datasets, this method can also be used on genomic reads from single individuals (Ågren et al. 2014). For this purpose we used a library of TE sequences based on several Brassicaceae species (Slotte et al. 2013). We used the default pipeline for PoPoolationTE, modified to require a minimum of 5 reads to call a TE insertion, and the procedure in Ågren et al. (2014) to determine heterozygosity or homozygosity of TE insertions. Parental origins of TE insertions were inferred by combining information from runs on F1s and their *C. rubella* parents. We used chi-square tests to assess tested whether the composition of heterozygous TE insertions targeted by uniquely mapping siRNAs differed from those not targeted by siRNAs.

We tested whether heterozygous TE insertions within a range of different window sizes close to genes (200 bp, 1 kbp, 2 kbp, 5 kbp, and 10 kbp) were associated with ASE by performing Fisher exact tests. We tested whether the expression of the allele on the same chromosome as a nearby (within 1 kbp) TE insertion was reduced compared to ASE at against genes without nearby TE insertions using a Wilcoxon rank sum test. Similar tests were conducted to test for an effect on relative ASE of TE insertions with uniquely mapping siRNAs.

## Acknowledgements

The authors thank Daniel Skelly, Duke University, for helpful advice on ASE analyses, Emily Josephs, University of Toronto and Adrian Platts, McGill University for information on conserved noncoding regions in *Capsella*, and Michael Nowak, Stockholm University, for valuable comments on the manuscript. Sequencing was performed by the SNP&SEQ Technology Platform in Uppsala. The facility is part of the National Genomics Infrastructure (NGI) Sweden and Science for Life Laboratory. The SNP&SEQ Platform is also supported by the Swedish Research Council and the Knut and Alice Wallenberg Foundation. The computations were performed on resources provided by SNIC through Uppsala Multidisciplinary Center for Advanced Computational Science (UPPMAX) under Project b2012122. This work was supported by grants from the Swedish Research Council, the Erik Philip-Sörensen foundation, the Nilsson-Ehle foundation, the Magnus Bergvall foundation, and the Royal Swedish Academy of Sciences to T.S. D.K. acknowledges funding from the Human Frontier Science Program (LT000783) and the German Research Foundation Priority Program 1529 – ‘Adaptomics’ (WE 2897).

## Supporting Information

S1 Information: S1 Text containing detailed procedures for filtering genomic regions, qPCR details, Supporting Figures (S2-S7) and Tables (S8-S16).

